# Molecular Docking of *Glycyrrhiza glabra* against the Conserved Target M1, NA and NS1 Proteins of Influenza A Viral Strains Identified through Pangenome Analysis

**DOI:** 10.1101/831461

**Authors:** S Aishwarya, Evangeline Shantha, E Nantha Devi, R Sagaya Jansi

## Abstract

Influenza viruses that infect humans are known to swiftly evolve over time. Influenza A virus has a negative single-stranded RNA genome in eight segments. Pangenome analysis of twelve strains of Influenza A viruses H1N1, H1N2, H2N2, H3N2, H5N1, H5N6, H7N2, H7N3, H7N7, H7N9, H9N2, and H10N8 gave insight on the core genes that are conserved and accessory genes that are specific for the strains. The proteins Neuraminidase, Matrix M1 and Nonstructural protein 1 were encoded by the core genes of segments 6, 7, and 8 respectively which proves that they are conserved in almost all the strains of influenza. The 3Dimensional structures of the core genes were interpreted by homology modeling and compared with corresponding Protein Data Bank structures (4MWQ, IEA3, 2GX9). Among several anti-viral phytocompounds that were virtually screened against the modeled and PDB target proteins, three molecules of Indian plant *Glycyrrhiza glabra* had high scores and interactions. Compounds 2,4,4’ Trihydrochalcone, Davidigenin and Licoflavone B docked well with the Neuraminidase, Matrix protein M1 and Nonstructural Protein NS1 respectively with good scores, minimized energy and interacted with the active sites. The compounds obeyed Lipinski’s Rule of five and exhibited drugability as well. Thus the present study focused on the drugable lead compounds from *glabra* that has inhibitory activities against the viral attachment, replication and matrix structure.

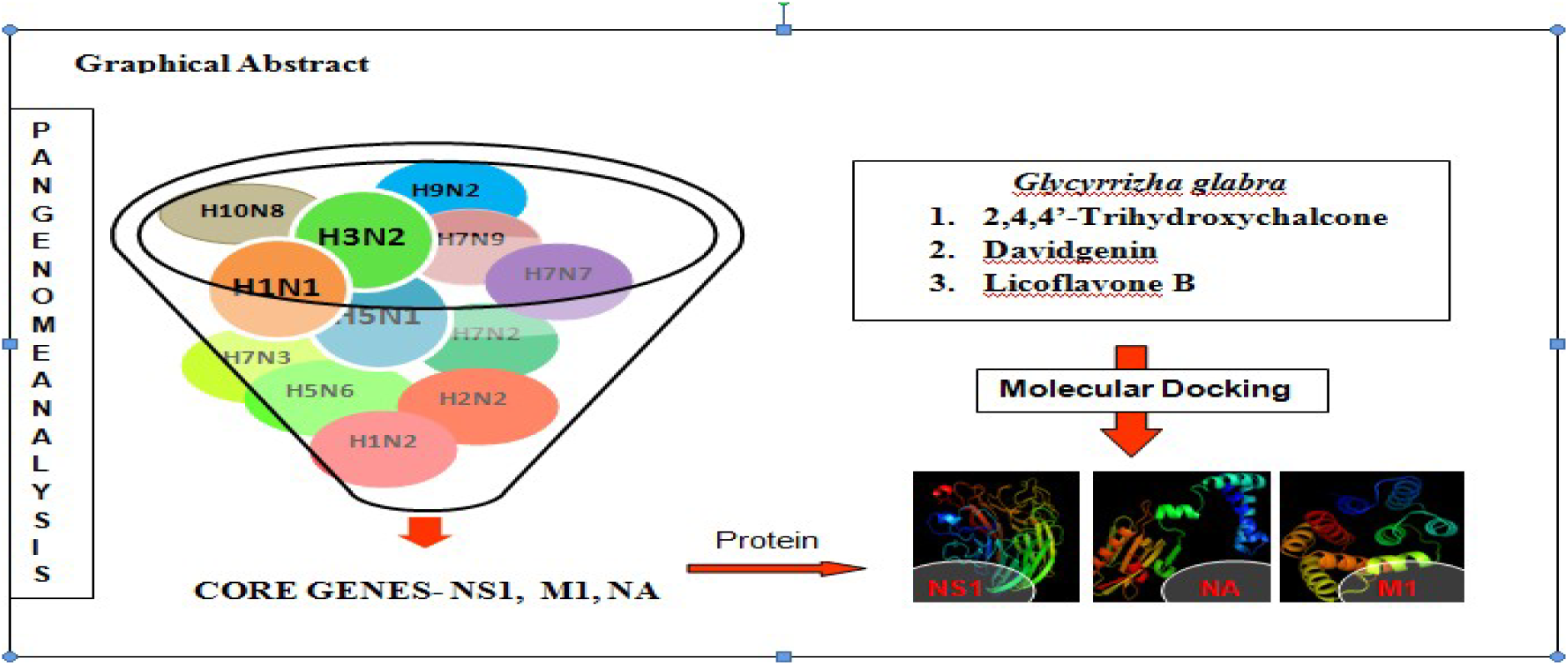

## Introduction

The most common cause of respiratory infections in humans is due to seasonal or epidemic outbreak of influenza viruses and recently there are many different strains identified (Aishwarya *et. al*., 2019). Influenza viruses belonging to the family *Orthomyxoviridae* are enveloped, with negative-single-stranded RNA genomes found in seven to eight gene segments (Miller, MS *et. al* 2013). Among the types A, B and C, Influenza A is more predominant in affecting humans and is mutated to form multiple strains. Influenza A viruses are subdivided by antigenic characterization of the Hemagglutinin (HA) and Neuraminidase (NA), a surface glycoprotein and occur in eighteen HA (H1–H16) and eleven NA (N1–9) subtypes (Hay AJ *et. al*, 2001). These subtypes are further divided into two or three groups: group 1 HA (H1, H2, H5, H6, H8, H9, H11, H12, H13, H16, H17, and H18) and group 2 HA (H3, H4, H7, H10, H14, and H15), or group 1 NA (N1, N4, N5, and N8), group 2 NA (N2, N3, N6, N7, and N9) and group 3 NA (N10 and N11) (Zhang *et. al*.,2014). Currently available treatment options like vaccines and anti-influenza drugs have a short protection plan and they become less effective over time due to the aberrant mutation rates of the viruses (ShinJ *et. al*., 2019).

Pangenome analysis of viruses enabled the identification of core genes that are conserved in multiple strains and enables identification of the conserved targets from the core genes (Darricarrere *et. al*, 2018). The conserved targets identified were Nonstructural protein 1 that is involved in viral-host interaction (Mok W *et. al*, 201), Matrix protein M1 is responsible for the viral envelope structure (Couch RB *et. al* 2014) and Neuraminidase enhances viral replication (Benton *et. al*, 2012). Though there are neuraminidase inhibitors available, their efficacy is debated since there are varied NA groups in different strains of the virus. Molecular docking is a computational approach that had accelerated drug design research with the efficient prediction of lead compounds against the 3-dimensional protein targets (Shanti N.,2011)

The current study aimed at identifying the core genes of multiple strains of influenza and determining the protein targets which were then studied for rigid molecular docking with the available antiviral plant compounds.

## Methodology

### Retrieval of complete genomes

The complete genomes of 12 different strains of Influenza A virus infecting humans were retrieved from NCBI Genome sequence portal (www.ncbi.nlm.nih.gov/)

### Pan-genome analysis

Pan-genome analysis of the retrieved segmented genes was performed using panseq (https://lfz.corefacility.ca/panseq/) to predict the core genes of the eight segments. The open reading frame was identified from the core genes and the translated protein segments were validated using BLAST –Basic Local Alignment Search Tool (www.blast.ncbi.nlm.nih.gov/).

### Comparative modeling of the translated protein sequences

Comparative modeling also called homology modeling is the best way to predict the structures of the proteins. The translated core genes of segments 6, 7 and 8 were modeled with suitable templates using the web server Phyre2 (http://www.sbg.bio.ic.ac.uk/phyre2).

### Library preparation

The entry point for any chemistry program within drug discovery research is generally the identification of specifically acting low-molecular-weight modulators with an adequate activity in a suitable target assay. Around 657 antiviral compounds were identified and retrieved from IMPPAT database. The library thus generated was optimized with an OPLS3 force field and stereoisomers were predicted using the ligprep module of Schr□dinger software.

### Screening of the druggable molecules

The prepared library compounds were screened based on the Lipinski’s rule of 5 that describes the drugability of the molecules based on the following four criteria

1. The molecular weight of the molecule is less than or equal to 500 Daltons
2. LogP (Ratio of Octanol to Water) is less than or equal to 5
3. Number of Hydrogen bond donor is less than or equal to 5
4. Number of Hydrogen bond acceptors is less than or equal to 10

### Target structure retrieval from PDB

Identifying and selecting the most appropriate drug target or receptor is the initial step in the drug designing procedure. Mostly proteins act as good targets for the drugs but in some cases, enzymes can also serve as excellent drug targets. Three viral targets namely Neuraminidase (PDB ID-4MWQ), Nonstructural protein NS-1 (PDB ID-2GX9) and Matrix protein M1 (PDB ID-1EA3) were selected and their 3D structure was retrieved from PDB (https://www.rcsb.org/) The available protein structures will enable the comparison of the modeled protein and its docking results.

### Protein Preparation and Receptor grid generation

The modeled proteins and the retrieved 3D protein structures were energy minimized and optimized using the Protein Preparation wizard of the Schr□dinger suite. The binding pocket of the modeled proteins was identified using the Computed Atlas of Surface Topography of Proteins (CASTP server -http://sts.bioe.uic.edu/castp/) and the all the six structures were subjected to grid generation through the receptor grid generation package of Schr□dinger suite for drug discovery.

### Molecular docking and interpretation of results

The prepared and filtered library compounds were subjected to rigid docking against the three sets of target protein structures using the Ligand Docking module of Schr□dinger. Standard precision mode screened out better-docked compounds and the next Xtra Precision mode yielded the best-docked compounds. The results were interpreted based on the minimized energy, docking score, and interacting residues

## Results and Discussion

### Retrieval of complete genomes

There were 12 different strains of Influenza A viral sequences available in the NCBI that were identified and sequenced at different time scales and periods. The complete genome sequences for the entire study were retrieved from NCBI and their accession numbers are listed in the table 1a and 1b.

**Table 1.** a List of Genomes of Influenza A strains b List of Genomes of Influenza A strains

| Year | 2016 | 2009 | 2018 | 2013 | 2012 | 1962 |
| --- | --- | --- | --- | --- | --- | --- |
| Strains | H1N1 | H1N2 | H3N2 | H5N1 | H7N3 | H2N2 |
| Segment 1 | KY509794 | JQ340008 | MH701027 | KF918495 | CY125725 | AY209940 |
| Segment 2 | KY509798 | JQ340007 | MH701026 | KF918496 | CY125726 | AY210014 |
| Segment 3 | KY509802 | JQ340006 | MH701025 | KF918497 | CY125727 | AY209996 |
| Segment 4 | KY509806 | JQ340003 | MH701020 | KF918498 | CY125728 | AY209959 |
| Segment 5 | KY509811 | JQ340005 | MH701023 | KF918499 | CY125729 | AY210079 |
| Segment 6 | KY509814 | JQ340004 | MH701022 | KF918500 | CY125730 | AY209908 |
| Segment 7 | KY509818 | JQ340009 | MH701021 | KF918501 | CY125731 | AY210037 |
| Segment 8 | KY509822 | JQ340010 | MH701024 | KF918502 | CY125732 | AY210164 |

**Table 1.b List of Genomes of Influenza A strains**
| Year | 2015 | 5-p 2003 | 2013 | 2017 | 2017 | 2013 |
| --- | --- | --- | --- | --- | --- | --- |
| Strains | H5N6 | H7N2 | H7N7 | H7N9 | H9N2 | H10N8 |
| Segment 1 | KT245150 | EU783920 | KF918334 | MF455313 | MF440743 | KJ406534 |
| Segment 2 | KT245149 | EU587369 | KF918335 | MF455314 | MF440742 | KJ406538 |
| Segment 3 | KT245148 | EU587370 | KF918336 | MF455315 | MF440741 | KJ406542 |
| Segment 4 | KT245143 | EU587368 | KF918337 | MF455316 | MF440736 | KJ406546 |
| Segment 5 | KT245146 | EU587371 | KF918338 | MF455317 | MF440739 | KJ406550 |
| Segment 6 | KT245145 | EU587372 | KF918339 | MF455318 | MF440738 | KJ406554 |
| Segment 7 | KT245144 | EU587373 | KF918340 | MF455319 | MF440737 | KJ406558 |
| Segment 8 | KT245147 | EU587374 | KF918341 | MF455320 | MF440740 | KJ406562 |

### Pangenome analysis

Pan-Genome Analysis was performed for all the 12 different strains of Influenza A virus infecting humans using Panseq. There were multiple core fragments and the most predominant core genes with less mutation were selected. The ORFs of the resulted Core fragments were predicted using ORF Finder. Table 2 represents ORFs of segments 6, 7 and 8 by ORF finder.

**Table 2.** ORFs of Core Fragments

| Segment Number | Open Reading Frames | Strand | Start | Stop | Length | Translated Protein |
| --- | --- | --- | --- | --- | --- | --- |
| Segment 8 | ORF 1 | + | <1 | 714 | 237 | Nonstructural Protein NS1 |
|  | ORF 2 | - | 767 | 117 | 216 |  |
|  | ORF 3 | + | 518 | >838 | 106 |  |
| Segment 7 | ORF 1 | + | <1 | 759 | 252 | Matrix Protein M1 |
|  | ORF 4 | + | 692 | >982 | 96 |  |
|  | ORF 11 | - | 278 | >3 | 91 |  |
|  | ORF 5 | - | 958 | 752 | 68 |  |
|  | ORF 9 | - | 330 | 157 | 57 |  |
| Segment 6 | ORF 2 | + | 221 | >910 | 229 | Neuraminidase NA |
|  | ORF 3 | + | 54 | 218 | 54 |  |
|  | ORF 6 | + | 567 | 692 | 49 |  |
|  | ORF 7 | - | 270 | 157 | 37 |  |

### Validation of the core genes

The validation of the Predicted core fragments was performed using Pfam sequence search. The segments 6,7 and 8 coded for the proteins neuraminidase, Matrix M1 and Nonstructural protein NS1 respectively and are represented in the figures 2.a, 2.b, and 2.c. Pfam results thus validated the protein-coding core genes of all 12 strains.

**Fig 1.**
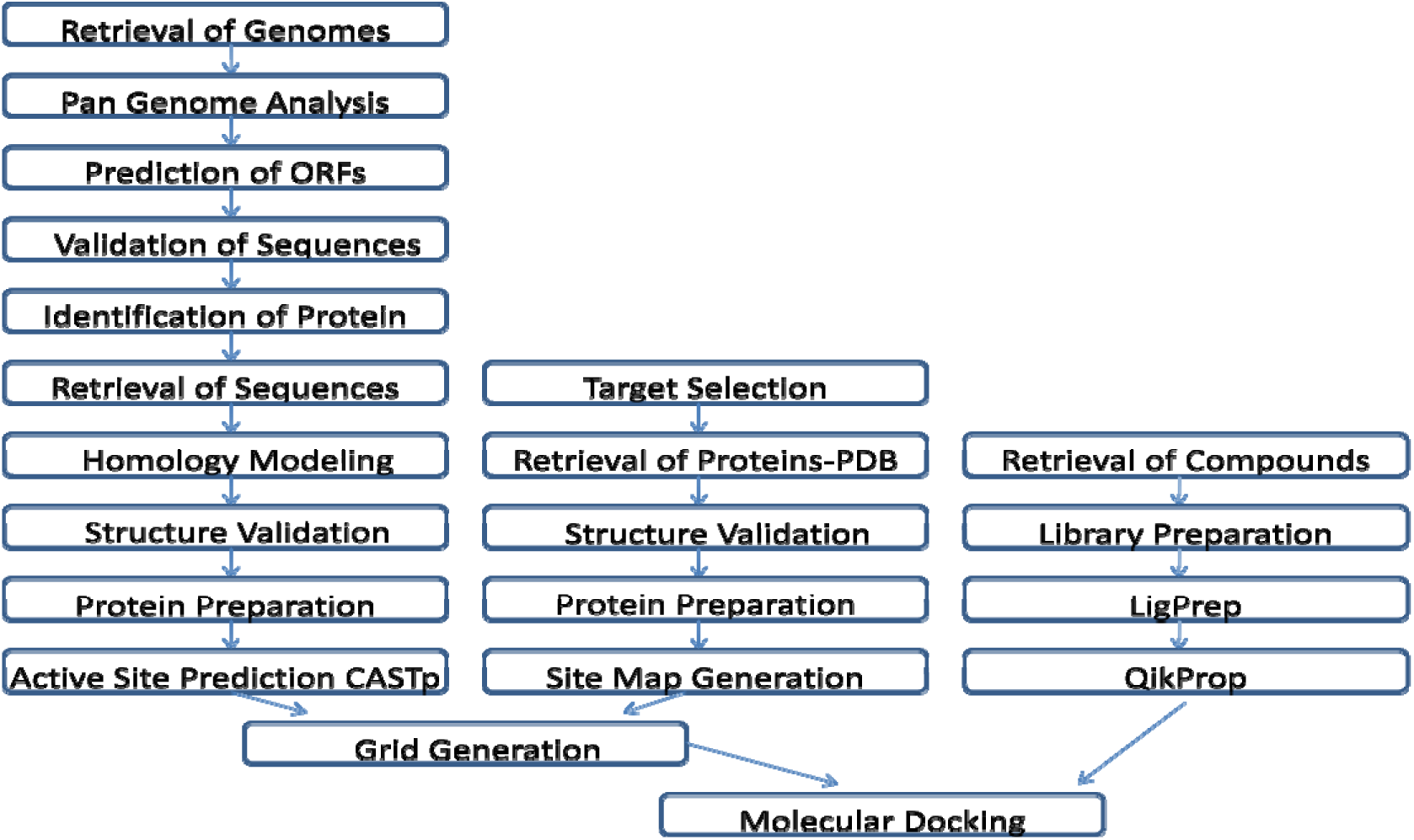
Flowchart represents the exact protocol that was followed in the current study

**Fig 2.**
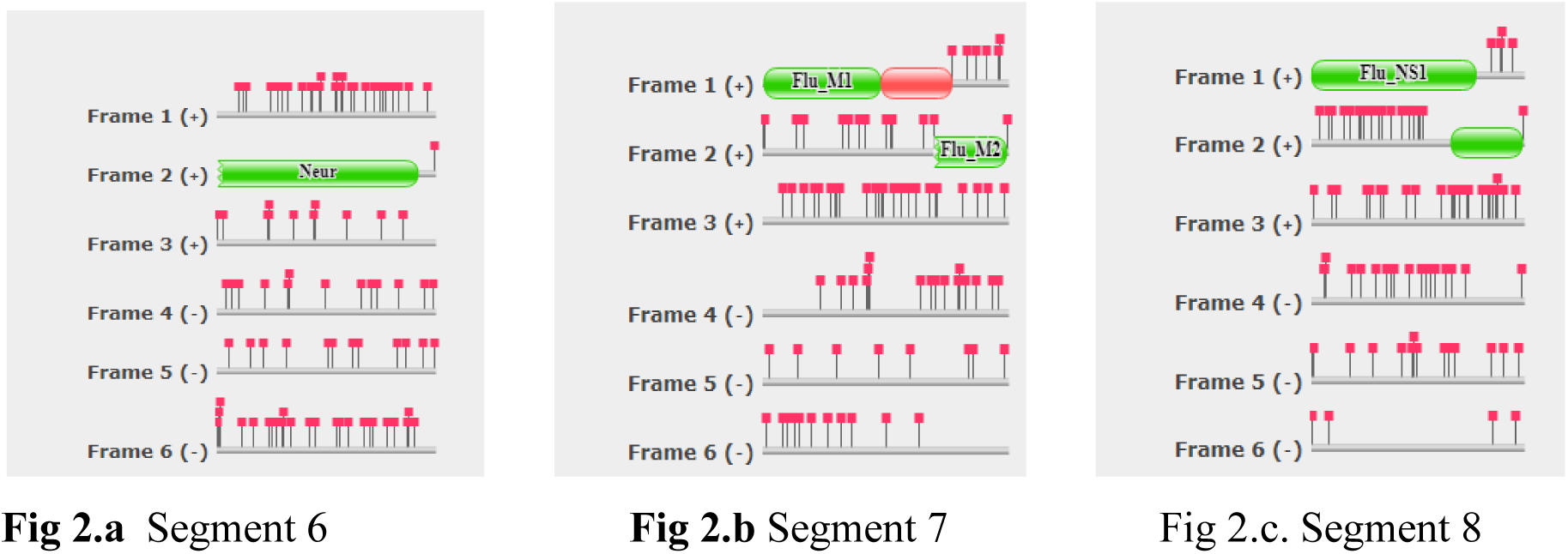

### Homology modeling

The ORF sequences were subjected to Blast against a nonredundant Protein database to identify the most similar sequence that was chosen as a template. The sequences that shared 100% identity and maximum query coverage was modeled comparatively and a 3 D structure of each protein was obtained.

The sequences were modeled using Phyre2 and models were generated. Predicted models were validated using rampage and the best model was retained for each segment and are represented in the figures 3.a, 3.b, and 3.c. Figures 4a, 4b and 4c represent the Ramachandran plot for the modeled proteins M1, NS1 and Neuraminidase respectively. The favored and allowed regions of residues are represented in table 4. The results thus show that the modeled protein structures are valid to be used for docking.

**Figure 3.**
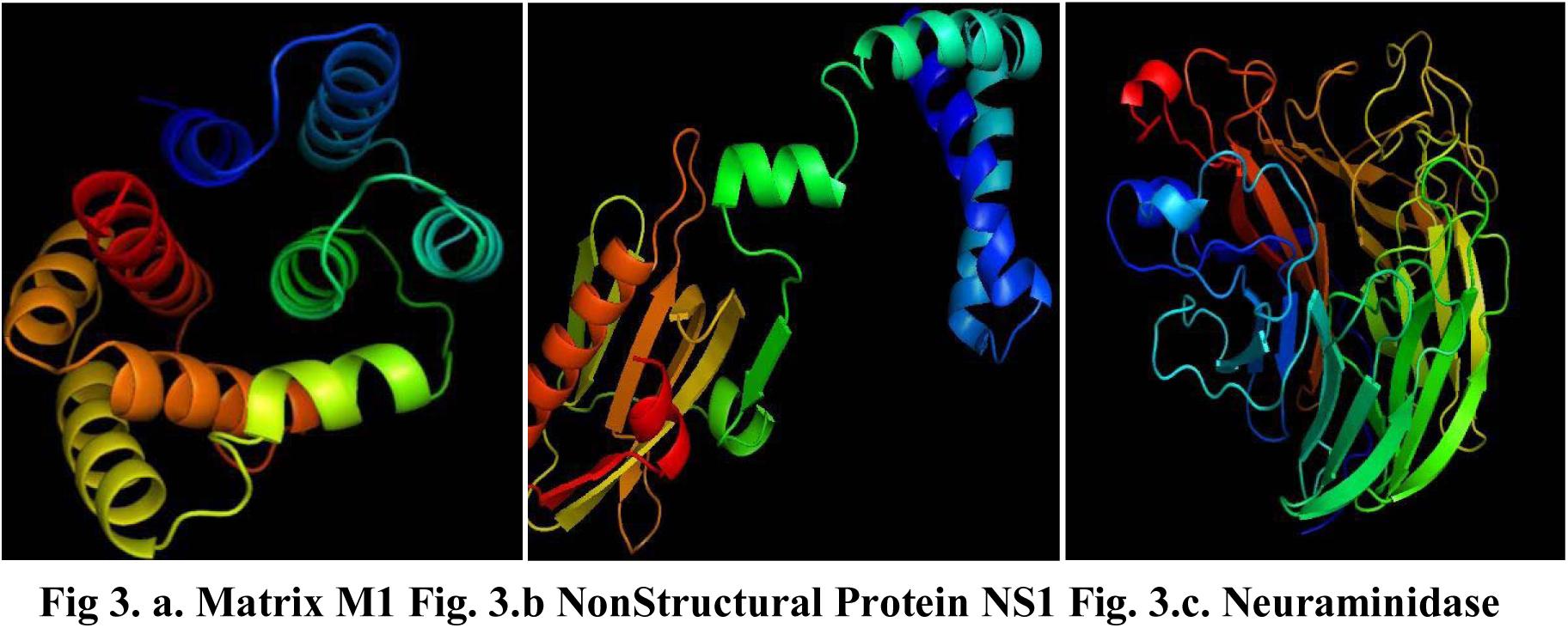
a −3.c Phyre2 Modeled proteins of the segments 6,7 and 8

**Figure 4.**
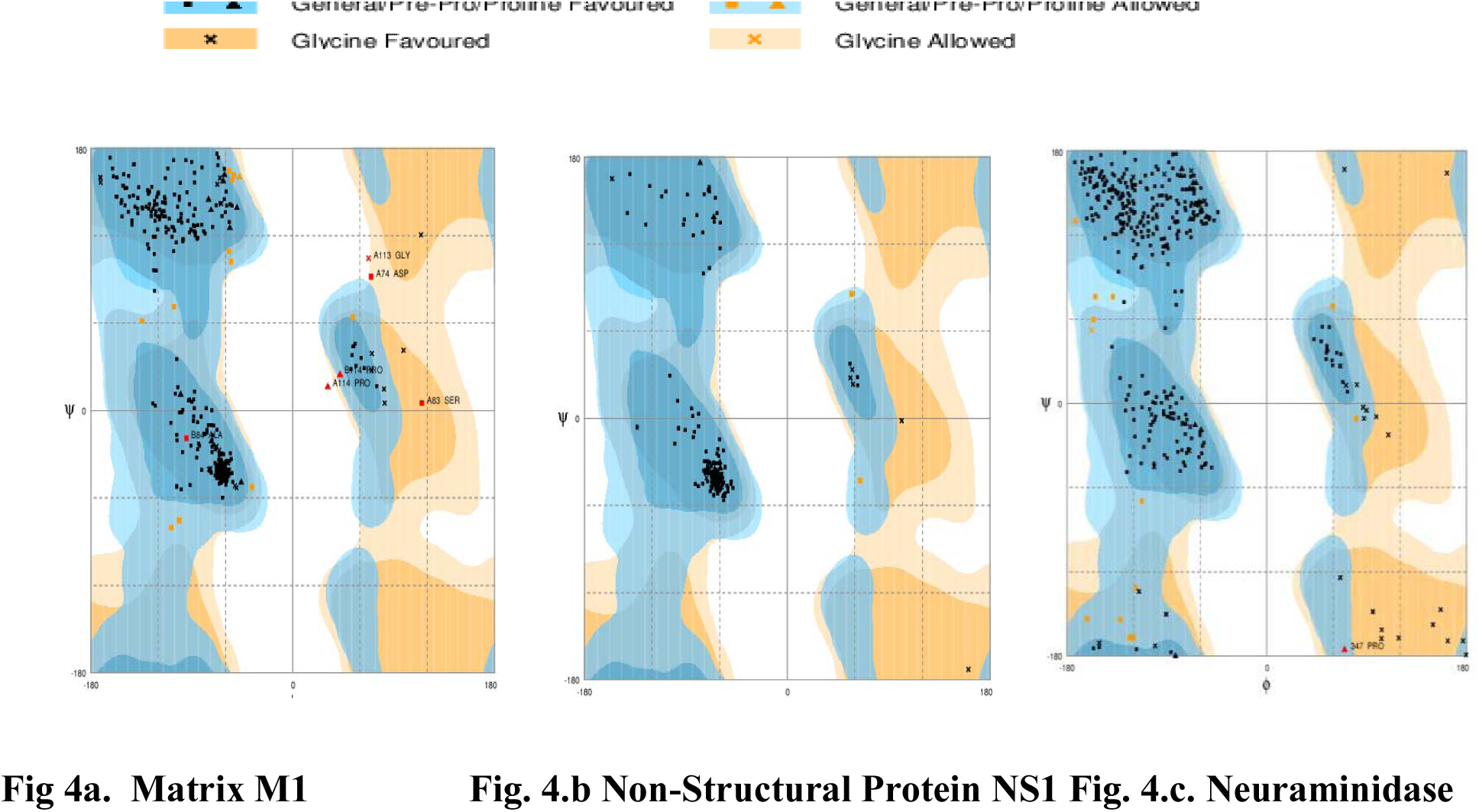
a – 4.c Ramachandran plots for validating the modeled protein

**Table 3.** BLAST results of the ORFs of segment 6,7 and 8

| Protein Name | Maximum score | Total score | Query coverage | E value | Identity % | UniProt ID |
| --- | --- | --- | --- | --- | --- | --- |
| Neuraminidase | 467 | 467 | 99% | 2.00E-165 | 98.69% | Q6XUA7 |
| Matrix protein 1 | 524 | 524 | 100% | 0 | 100.00% | Q6XT42 |
| *Nonstructural* protein 1 | 478 | 478 | 100% | 6.00E-173 | 97.47% | P69277 |

**Table 4.** Results of Ramachandran plot for the three modeled proteins

| Protein | # residues in the Favoured region | # Residues in the Allowed region | # Residues in the Outlier region |
| --- | --- | --- | --- |
| Matrix Protein 1 | 154 (98.7%) | 2 (1.3%) | 0 (0.0%) |
| *Nonstructural* Protein 1 | 378 (95.5%) | 12 (3.0%) | 6 (1.5%) |
| Neuraminidase | 372 (96.4%) | 13 (3.4%) | 1 (0.3%) |

### Retrieval of structures from PDB

The known structures of the proteins coding the segment 6, 7 and 8 were retrieved from Protein Data Bank based on the resolution. The Ramachandran plot analysis was performed for the retrieved protein structures whose values are mentioned in table 5.

**Table 5.** Results of Ramachandran Plot for the three proteins retrieved from PDB

| Protein | PDB ID | # residues in the Favoured region | # residues in the Allowed region | #residues in the Outlier region |
| --- | --- | --- | --- | --- |
| Matrix Protein 1 | 1EA3 | 269 (86.8%) | 26 (8.4%) | 15 (4.8%) |
| NonStructural Protein 1 | 2GX9 | 237 (96.7%) | 8 (3.3%) | 0 (0.0%) |
| Neuraminidase | 4MWQ | 372 (96.4%) | 14 (3.6%) | 0 (0.0%) |

### Active site prediction

The prediction of Active site was performed using the CASTp server for modeled protein and for the PDB proteins using sitemap module of Schr□dinger. The number of binding sites and the residues involved are represented in table 6.

**Table 6.** Active site residues predicted using CastP web server

| Protein | Binding site for modeled protein using Castp |  | Binding site for PDB proteins using Sitemap |  |
| --- | --- | --- | --- | --- |
|  | # Binding Sites | Residues | # Binding Sites | Residues |
| Matrix Protein 1 | 1 | 81, 84, 85, 93, 96, 126, 130, 133, 135 | 1 | 13-16, 47-57, 79, 82-89, 92, 93, 121, 122, 125, 126, 154, 157, 158 |
| Nonstructural Protein 1 | 1 | 13, 17, 20, 21, 24, 44, 58, 62, 66 | 1 | 89, 91, 92, 94-97, 99, 131-133, 145-148, 150 |
| Neuraminidase | 1 | 118, 119, 134, 151, 152, 156, 178, 179, 221, 222, 224, 225, 227, 240-242, 246, 252, 275-278, 292, 371, 406 | 1 | 120, 135, 152, 153, 157, 180, 181, 223, 224, 226-229, 242-244, 246-248, 277-280, 294, 296, 406 |

### Ligand Preparation

Around 19 anti-viral plants of Indian origin were selected and their respective 657 phyto-compounds were identified and their 2D structures were retrieved from Indian Medicinal Plant, Phytochemistry, and Therapeutics (IMPPAT database-https://cb.imsc.res.in/imppat/) through PubChem. The compounds were optimized and energy minimized using the LigPrep module of Schr□dinger Suite.

### Virtual Screening of Ligands

The compounds were filtered based on properties that include molecular weight, hydrogen donors, hydrogen acceptors, QPlogKp values, Percentage of Human oral Absorption using QikProp. Around 363 Ligands were filtered out based on the said parameters and were used for molecular docking.

### Molecular Docking

Rigid molecular docking was carried out with the virtually screened phytocompounds and the targets that were processed. The results of the docking score, energy and interaction with residues were interpreted in both SP and XP modes. The docked results of Neuraminidase, Matrix M1 and Non-Structural Protein NS1 are represented in tables 7, 8 and 9 respectively.

**Table 7:**
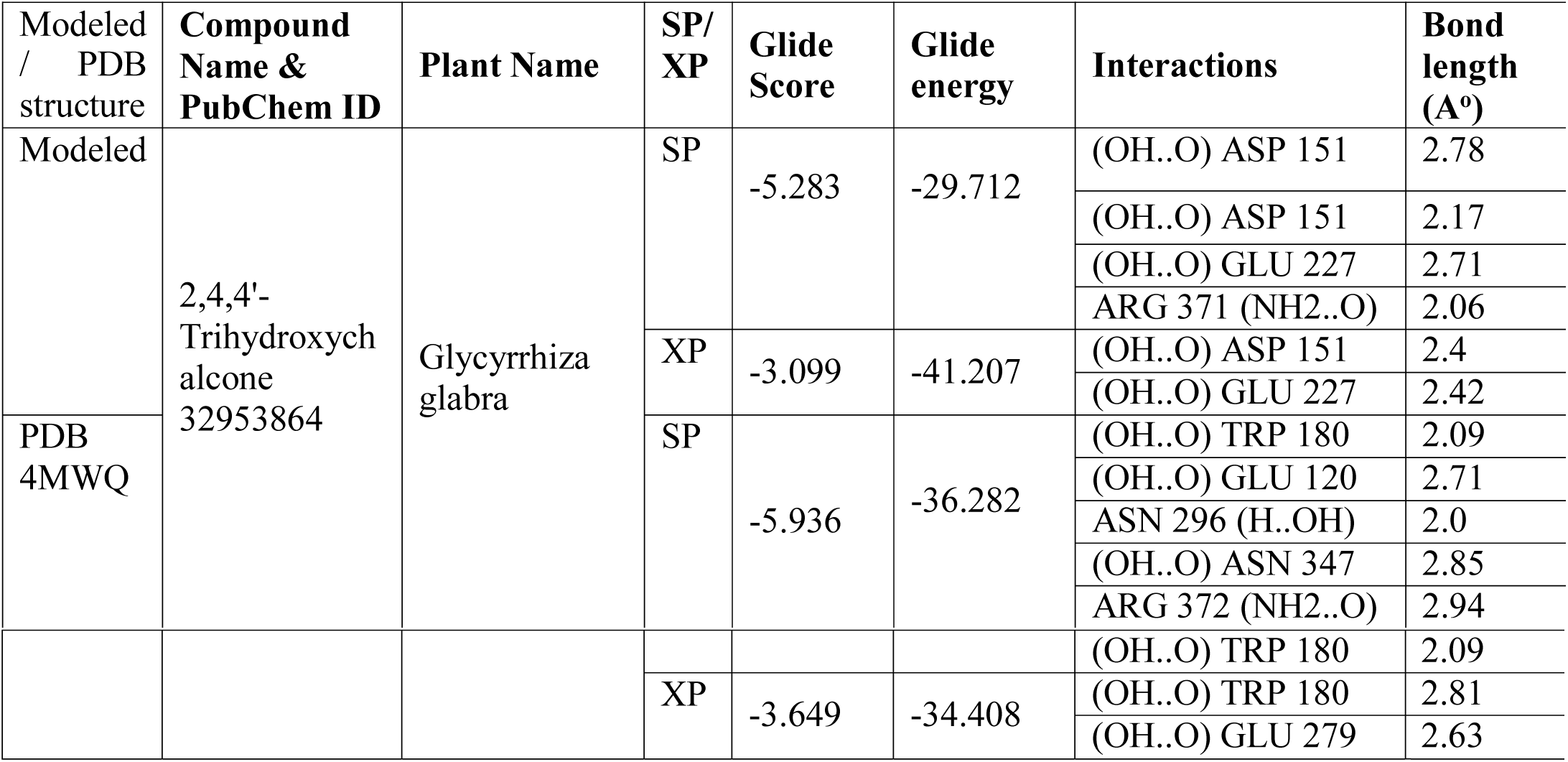
Docking Results of the Neuraminidase protein

**Table 8.** Docking results of Matrix protein

| Modeled / PDB structure | Compound Name & PubChem ID | Plant Name | SP/ XP | Glide Score | Glide energy | Interactions | Bond length (Å°) |
| --- | --- | --- | --- | --- | --- | --- | --- |
| Modeled | Davidigenin<br>442342 | Glycyrrhiza glabra | SP | -6.885 | -36.249 | (OH..O) SER 126 | 2.64 |
|  |  |  |  |  |  | (OH..O) ASN133 | 2.81 |
|  |  |  |  |  |  | (OH..O) MET135 | 2.92 |
|  |  |  | XP | -5.71 | -36.108 | (OH..O) SER 126 | 2.64 |
|  |  |  |  |  |  | (OH..O) MET 135 | 2.95 |
|  |  |  |  |  |  | (OH..O) ASN 133 | 2.8 |
| PDB IEA3 |  |  | SP | -6.012 | -33.074 | lig(OH..O) ASP 89 | 2.71 |
|  |  |  |  |  |  | lig(OH..O) ASN 85 | 2.21 |
|  |  |  | XP | -4.993 | -32.676 | (OH..O) ASN 85 | 2.12 |
|  |  |  |  |  |  | (OH..O) SER 157 | 2.9 |

**Table 9:**
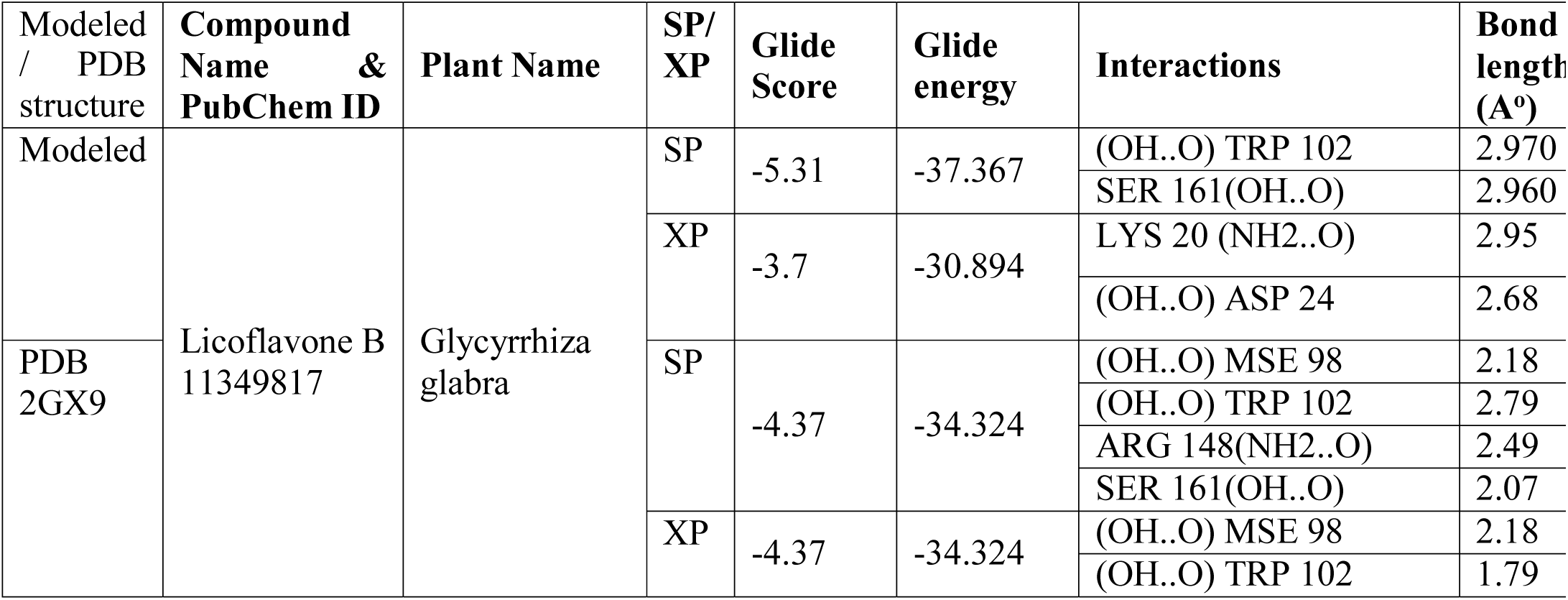
Docking results of the Neuraminidase protein

Figures 6, 7 and 8 show the docked images of the ligands in both SP and XP modes.

**Fig 5.**
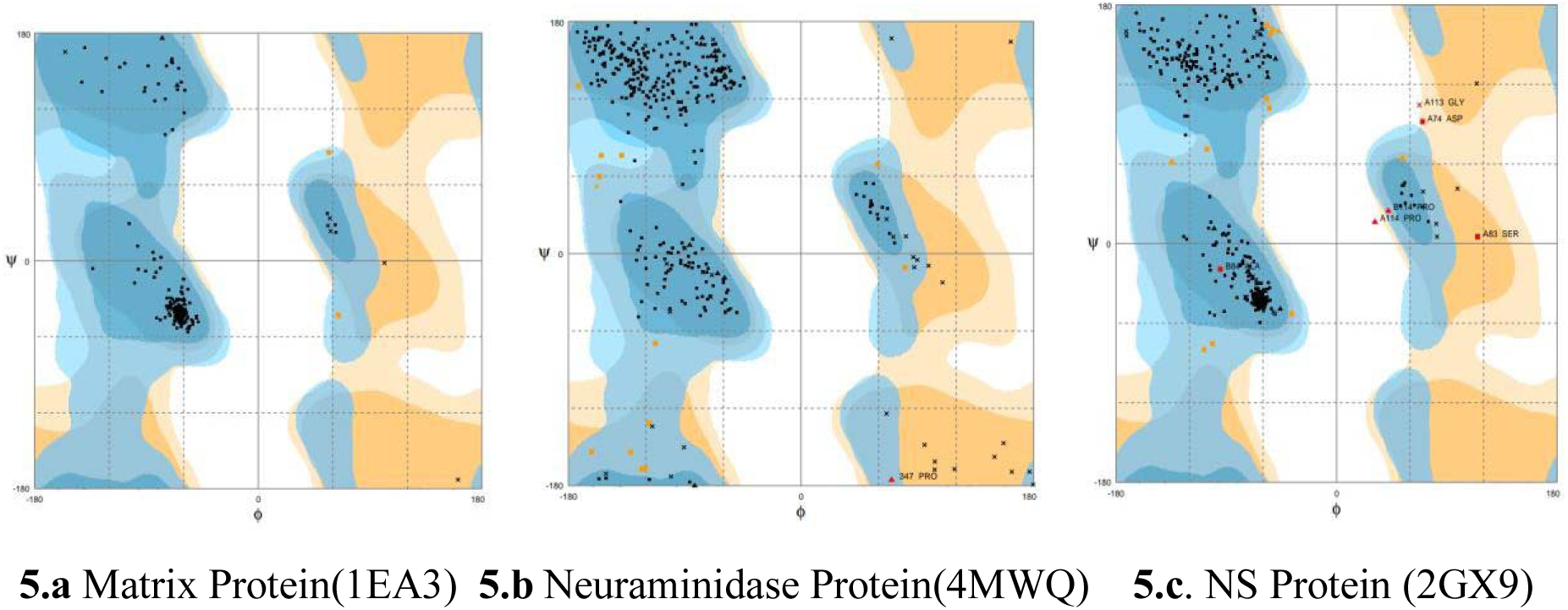
Ramachandran plot analysis for PDB retrieved proteins

**Fig 6.**
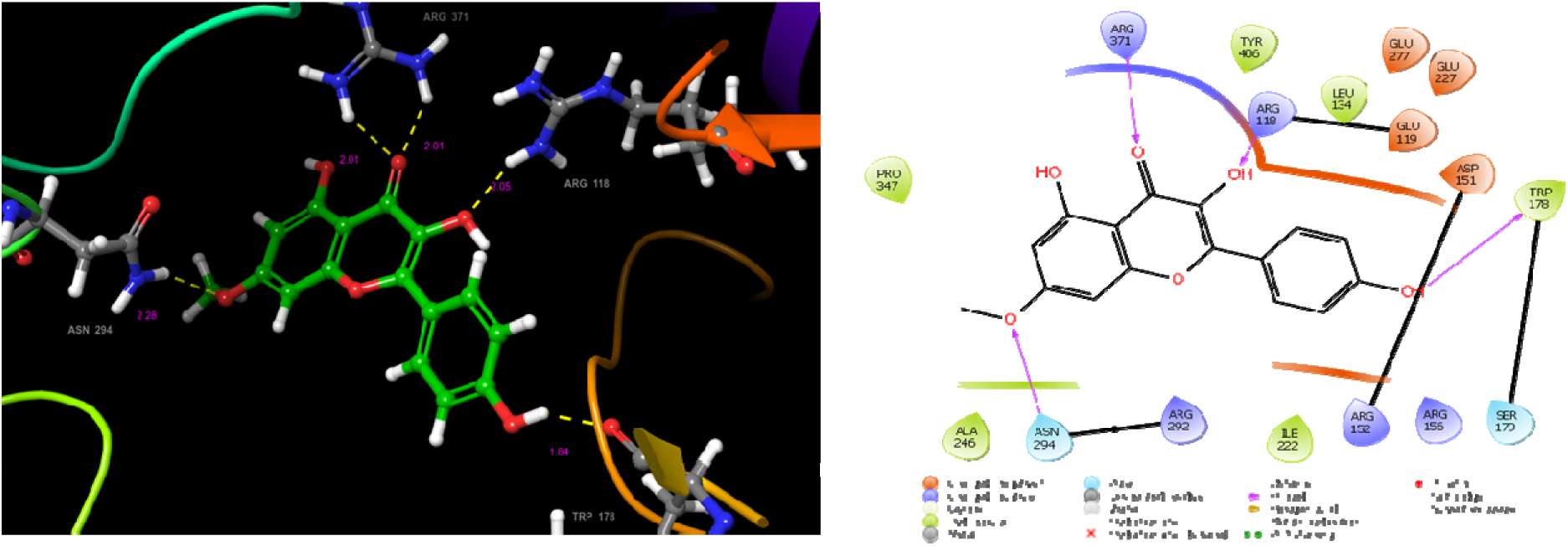
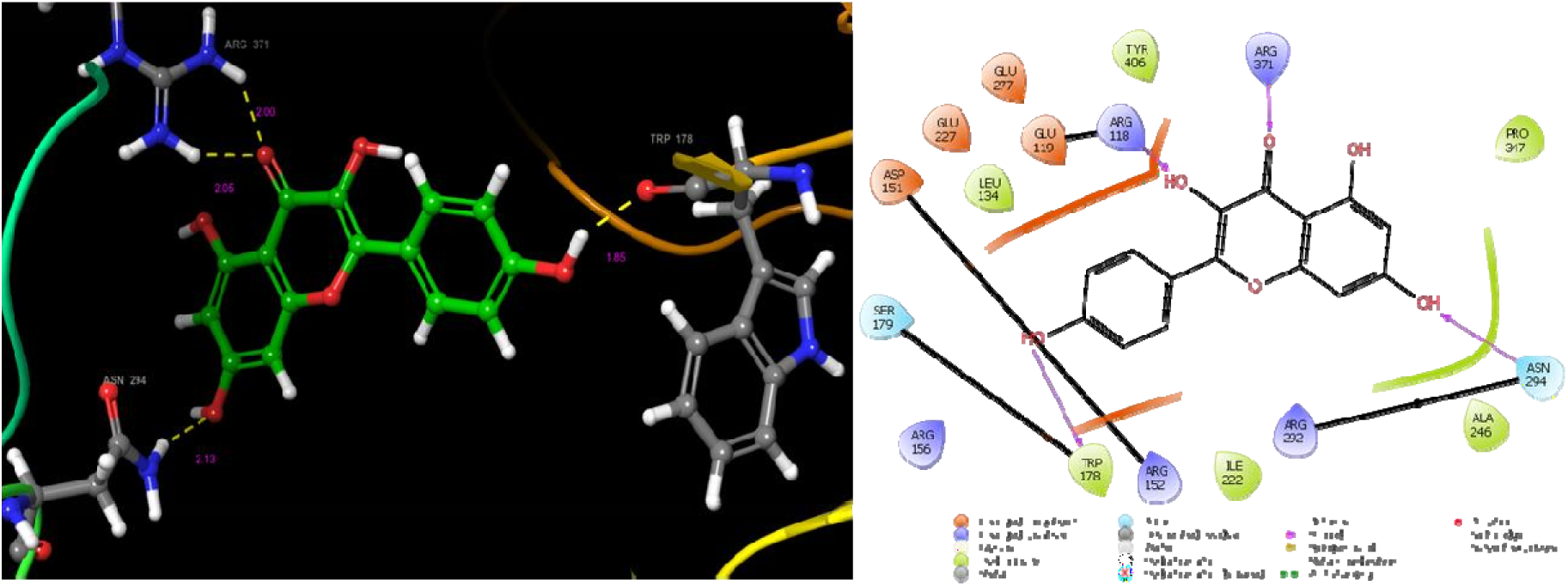
a: 2,4,4’ Trihydroxychalcone binding with the modeled neuraminidase protein 6 b: 2,4,4’ Trihydroxychalcone binding with the PDB structure of neuraminidase protein (4MWQ)

**Fig 7.**
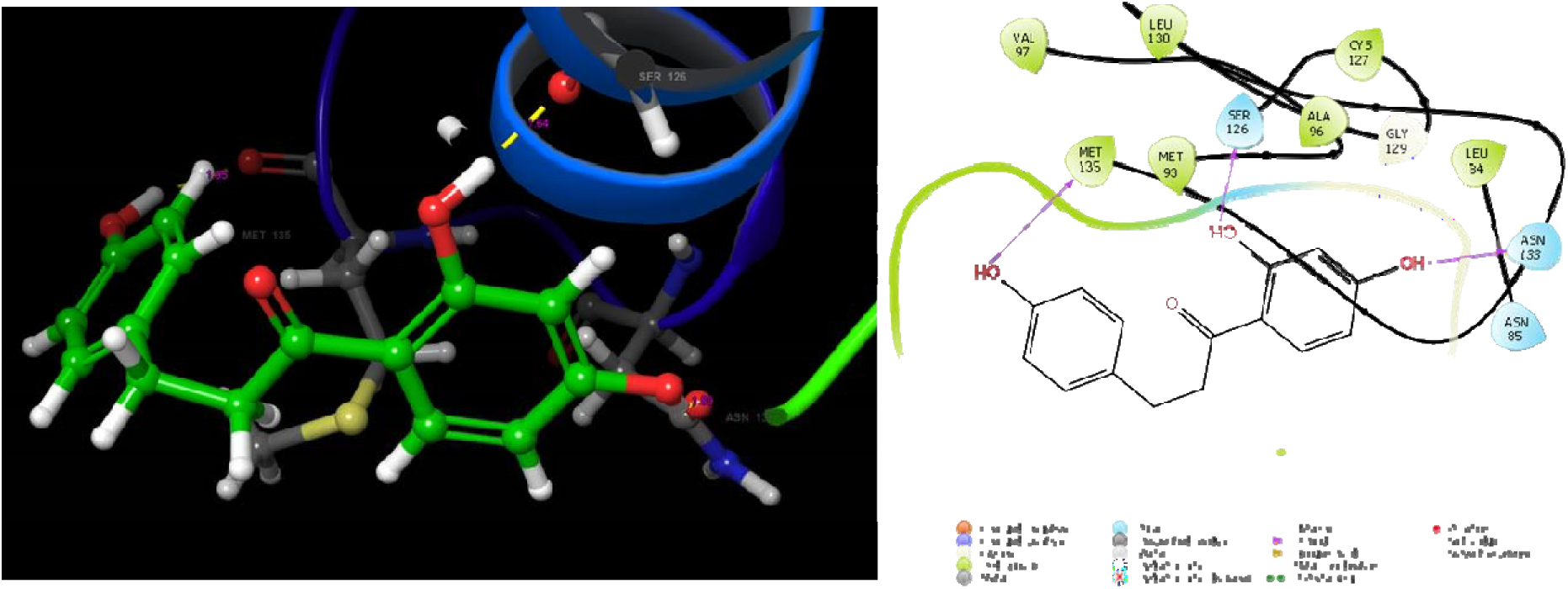
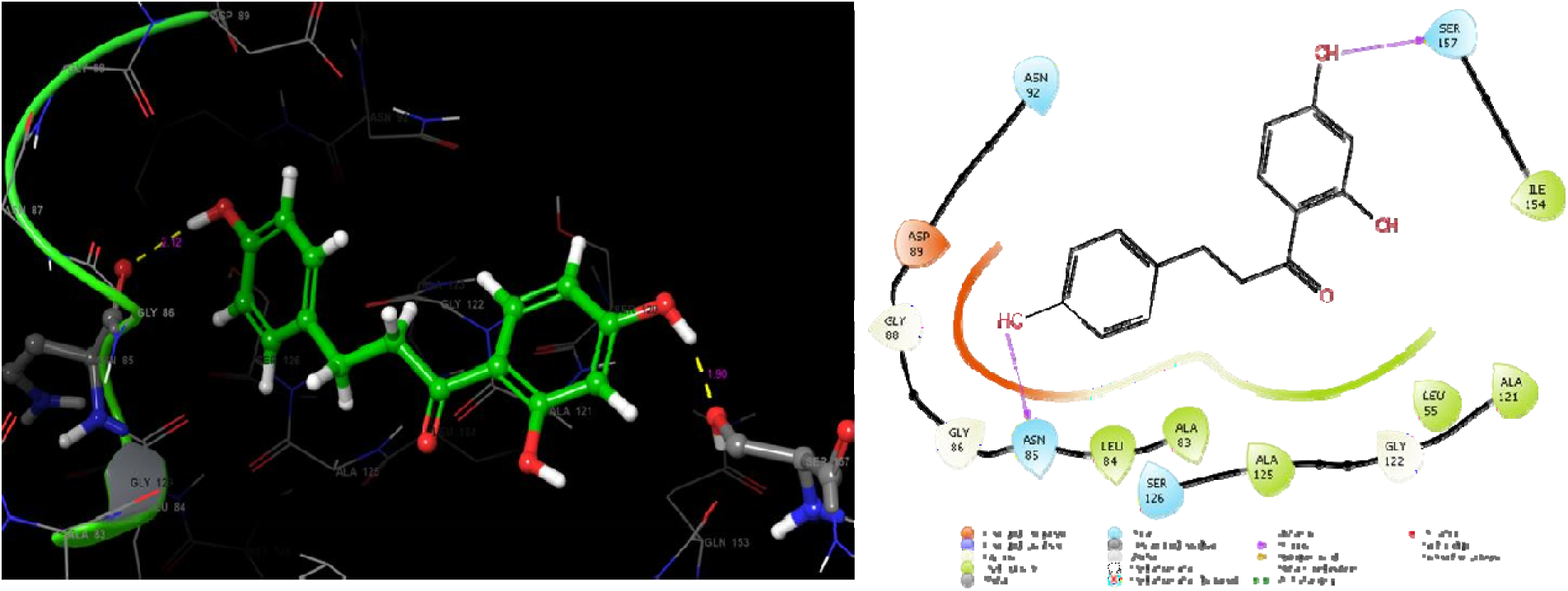
a Davidigenin docking with modeled Matrix M1 b Davidigenin docking with PDB Matrix M1 (IEA3)

**Fig 8.**
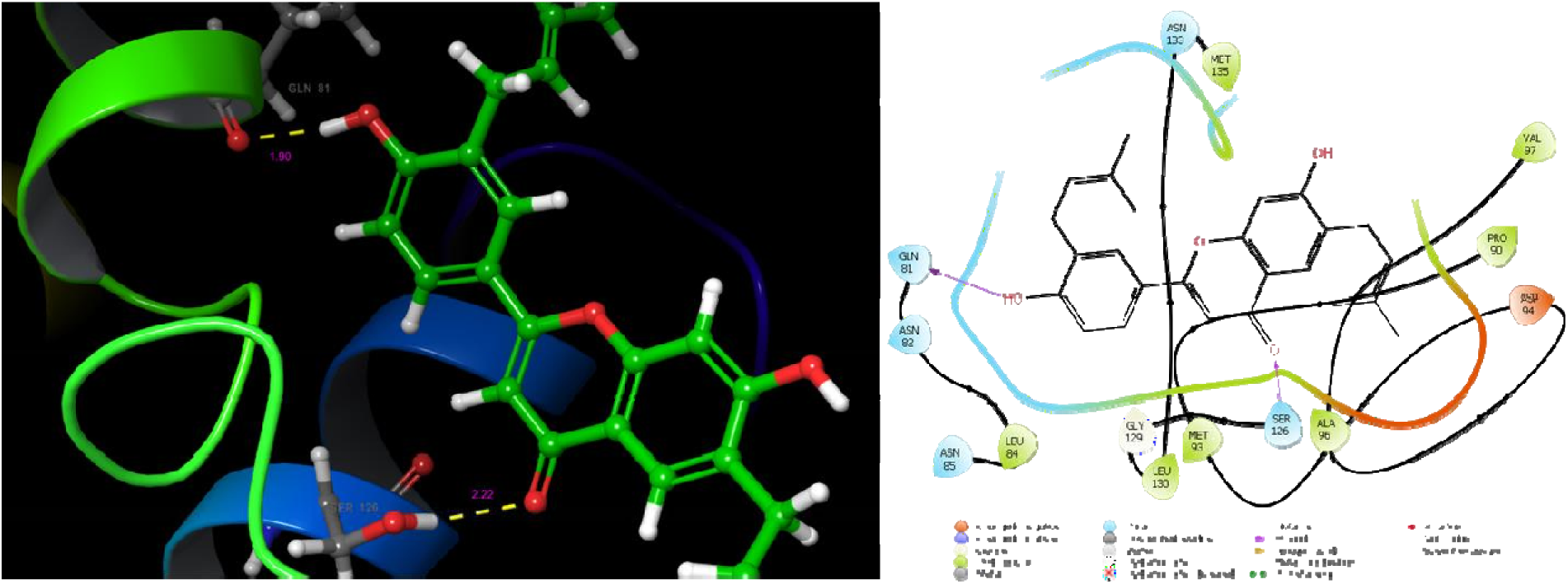
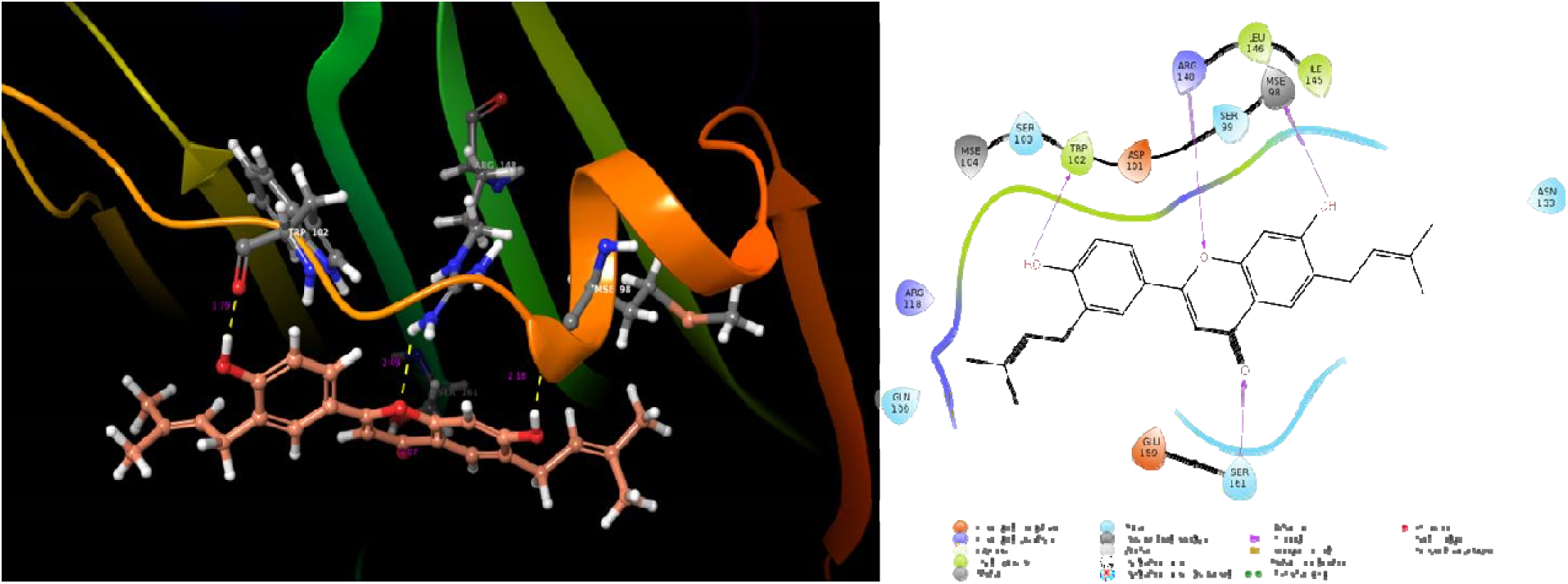
a Licoflavone B docked with Neuraminidase modeled protein b Licoflavone B docked with PDB Neuraminidase protein (2GX9)

Among the 392 virtually screened phytocompounds, 2,4,4’ Trihydroxychalcone from *Glycyrrhiza glabra* was found to exhibit good glide score and minimal energy against the neuraminidase protein. The interactions were also good with the active site residues of both the modeled protein and PDB structures. The compound also obeyed Lipinski rule of five with the molecular weight of 256.29g/mol, LogP value of 2.6, 3 Hydrogen bond donors and 4 hydrogen bond acceptors. Percentage of human oral absorption was 92% which makes the compound an effective lead for drug development.

Among the 392 virtually screened phytocompounds, Davidigenin from *Glycyrrhiza glabra* was found to exhibit good glide score and minimal energy against the matrix protein M1. The interactions were also good with the active site residues of both the modeled protein and PDB structures. The compound also obeyed Lipinski rule of five with the molecular weight of 258.23g/mol, LogP value of 3, with 3 Hydrogen bond donors and 4 hydrogen bond acceptors. Percentage of human oral absorption was 89% which makes the compound an effective lead for drug development.

Among the 392 virtually screened phytocompounds, Licoflavone B from *Glycyrrhiza glabra* was found to exhibit good glide score and minimal energy against the matrix protein M1. The interactions were also good with the active site residues of both the modeled protein and PDB structures. The compound also obeyed Lipinski rule of five with the molecular weight of 390.479g/mol, LogP value of 6.3, with 2 Hydrogen bond donors and 4 hydrogen bond acceptors. Percentage of human oral absorption was 88% which makes the compound an effective lead for drug development.

## Conclusion

The ideal way to find suitable targets using computational approaches for any particular study of interest has been found to be effective in recent years. Influenza A viral pandemic outbursts are more common and before a valid treatment is suggested there is always a huge loss of lives. There has been an ample rate of mutation taken place that has given rise to around 18 different strains. This study has paved way for finding the potential targets from the genomic level of whole genomes of 12 different strains of Influenza A virus which has been recorded to infect humans that included H1N1, H1N2, H3N3, H5N1, H7N3, H2N2, H5N6, H7N2, H7N7, H7N9, H9N2, H10N8. The Influenza virus is 8 segmented single-stranded negative RNA virus where each segment codes for a protein. The pan-genome analysis revealed that though the strains were continuously mutating over time, there are a set of genes and proteins that are strongly conserved. The segments 6, 7, and 8 were included in this study because they were less targeted. Virtually screening of antiviral compounds led to the identification of novel lead compounds, 2,4,4’ Trihydroxychalcone, davidigenin and licofalvone B from the well known Indian Plant *Glycyrrhiza glabra*. The current study is an extensive computational approach starting from comparative genome analysis, validating the core genes, identifying the acceptable targets and predicting leads that inhibit the targets. The study requires an investigational invitro and invivo tests done subsequently to be able to prove but still the results suggested here is valid and recommends proceeding for further research. If successful we will be able to target multiple strains of influenza viruses at the shortest time span.

## Acknowledgment

Deep gratitude to DST-FIST grants which had enabled us to procure the state of art drug development Schr□dinger suite.

## Abbreviations used

BLAST: Basic Local Alignment Search Tool
BLASTP: Basic Local Alignment Seach Tool of Proteins
ORF: Open Reading Frames
CASTp: Computed Atlas of Surface Topography of Proteins
ADME: Absorption, Distribution, Metabolism, and Excretion
SP: Standard Precision
XP: Extra Precision
PDB: Protein Data Bank

